# Inhibiting nociceptor endocytosis reduces MIA-induced osteoarthritic pain behavior

**DOI:** 10.64898/2026.08.21.746311

**Authors:** Aubrey J. Cooper, Jessica S. Tabman, Raider Rodriguez, Arin Bhattacharjee

**Affiliations:** Program for Neuroscience, University at Buffalo - The State University of New York, Buffalo, New York 14203, USA; Pharmacology and Toxicology, University at Buffalo - The State University of New York, Buffalo, New York 14203, USA

## Abstract

**Introduction:** Osteoarthritis (OA) is a degenerative joint condition characterized by chronic pain and the need for pain management. Locally targeting the endocytotic AP2 complex in nociceptors presents a potential strategy for providing sustained pain relief in individuals with OA.

**Objective:** We investigated whether pain behavior associated with OA can be mitigated by genetically silencing the AP2α2 subunit of the AP2 complex in nociceptors and by pharmacologically inhibiting the AP2 complex through the intraarticular administration of a small lipidated decoy peptide.

**Method:** Monoiodoacetate (MIA) was employed to induce knee joint OA in mice and rats. Pain behavior was assessed using dynamic weight-bearing and von Frey filaments. Upon confirmation of established OA pain behavior, in vivo AP2α2 genetic knockdown in mice was achieved through sciatic nerve transfection of a targeting AP2α2 short hairpin RNA (shRNA). To pharmacologically target endocytosis, a single intraarticular injection of peptide was administered into the arthritic knee of rats. The injection contained either the AP2 inhibitor peptide or a scrambled peptide control.

**Results:** Pain behavior was significantly reduced after both genetic and pharmacological disruption of AP2-driven endocytosis. Animals treated with the Ap2 inhibitor peptide exhibited reduced pain behavior throughout the 28-day assay period. Following the completion of behavioral testing, arthritic knee joints and contralateral healthy knee joints were subsequently collected to assess the impact of the treatment on disease progression. Micro-computed tomography analysis revealed a preservation of bone volume in the arthritic joints that received the AP2 inhibitor peptide treatment, in contrast to the scrambled peptide group.

**Conclusion:** These findings demonstrate that the inhibition of nociceptor endocytosis by a small lipidated peptide presents a promising approach to provide sustained relief from joint pain in individuals with arthritis.

## 1. Introduction

Osteoarthritis (OA) is a degenerative joint condition that leads to chronic pain and a need for pain relief. OA is characterized by inflammation, cartilage breakdown, early subchondral bone degradation followed by bone remodeling[8]. Conservative pharmacological approaches such as intraarticular injections of pharmacological therapies are used before resorting to joint replacement surgery. However, current intraarticular therapeutics are often inadequate in pain relief and disease control[23].

Ongoing internal pain, the type of pain that requires coping responses, are mediated by TRPV1-positive nociceptors[7]. Indeed, it has been argued that for patients with ongoing injury-associated pain and associated inflammation, targeting specifically the TRPV1/CGRP^+^ class of afferent fibers may be key in providing effective pain relief[14]. Synovial tissue happens to be densely innervated by TRPV1/CGRP^+^ expressing nociceptors[13; 21]. Additionally, mouse and human CGRP^+^ nociceptors preferentially express the endocytotic adaptor protein 2 (Ap2) complex subunit called Ap2α2[15], which is part of the multimeric AP2 complex[22]. This is one of two genes that encodes the α subunit of the AP2 complex[15]. During *in vivo* AP2α2 knockdown it was determined that nociceptor endocytosis was required for both the development and the maintenance of inflammatory pain[15]. A small AP2 decoy peptide that targets the AP2σ2 subunit[10], when lipidated, blocked Slack K_Na_ channel endocytosis and prevented dorsal root ganglion (DRG) neuronal hyperexcitability after protein kinase A stimulation[4]. Pharmacologically, this peptide provided sustained analgesia after a single administration (many days) in multiple recoverable inflammatory pain models[15].

Clinically effective intraarticular therapies for OA pain, however, require weeks of pain relief rather than days. Utilizing the monoiodoacetate (MIA) model of OA pain, we aimed to determine the efficacy of a small lipidated decoy peptide that targets the AP2σ2 subunit, designated as AP2σ inhibitor peptide, in mitigating OA pain behavior following intraarticular injection. Using micro-computed tomography (microCT) analysis, we also examined the effects of the AP2σ inhibitor peptide on OA-related bone pathophysiology.

## 2. Method

### 2.1 Animals

Male and female 250-275 g Sprague Dawley (SD) rats and male and female 8–10-week-old C57BL/6 mice were purchased from Envigo (Indianapolis, IN). Animal procedures, including injections, surgeries, and experiments were approved of by the Institutional Animal Care and Use Committee (IACUC) at The State University of New York at Buffalo. Formal National Institute of Health (NIH) guidelines were adhered to during all experiments. Upon arrival at the Laboratory Animal Facilities (LAF) all animals were allotted separate home cages, as was protocol approved. An acclimation period of seven days to allow animals time to recover from any stress induced during shipment was allotted before behavioral experiments commenced. All injections and pain monitoring assays occurred during the light phase. Animals received food and water ad libitum during a 12-hour light/dark cycle. The experimenter was blinded to all experimental conditions, with unblinding occurring only after the analysis.

### 2.2 Monoiodioacetate induction of arthritis

Osteoarthritis was induced by unilateral intraarticular injection of MIA (Sigma-Aldrich, St. Louis MO) as previously described[19]. Mice were anesthetized by 2% inhalation isoflurane and placed supine on a heating pad. Right knees were injected with 0.8mg MIA in 8μL of phosphate buffered saline (PBS) using a 30-guage X ½ inch needle on a Hamilton syringe (Hamilton, Reno NV). For rats, 2mg MIA in 30μL of PBS was injected using a 28-guage X ½ inch needle. Injection depth was kept at 2 mm for mice and 5 mm for rats using a polyethylene tube to prevent further needle penetration. To ensure MIA stayed within the joint cavity, the needle was maintained in the knee joint for at least 60s after injection to avoid backflow. Manual manipulation (bending the knee up and down) for 60 seconds after the needle was removed helped to coat the joints evenly.

### 2.3 Sciatic Nerve transfection of shRNAs

*In vivo* sciatic nerve transfection of shRNAs were performed as previously described [12; 15; 16; 19]. C57Bl/6 mice were anesthetized with 2% isoflurane. A 3 cm posterior longitudinal incision was made at the lumbar region of the spine. Sterile toothpicks were used to separate the paraspinal muscles exposing the sciatic nerve. The *in vivo* transfection reagent used was invivo-jetPEI®(Sartorius Bohemia, NY). 3μL of PEI/shRNA plasmid DNA polyplexes was directly injected into the sciatic nerve using a Hamilton syringe with a 32-guage blunt needle. A mouse specific Ap2α2 shRNA and control shRNA were purchased from Santa Cruz Biotechnology (Santa Cruz, CA).

Following injection, the needle was kept in the sciatic nerve for at least 1 minute to promote diffusion of the polyplexes. After injection, the incision was closed with wound clips and animals were observed until full recovery.

### 2.4 Peptides

The lipidated AP2σ targeting peptide used was as previously described[4; 15]. The sequence of the AP2σ peptide was RMS^P^<u>EIKRLL</u>SE, where the first serine was phosphorylated (designated by a superscript ‘P’). Underlined sequence designates the acidic di-leucine motif that binds to the Ap2σ2 subunit. The scrambled sequence was IERLSEMS^P^LRK. Both peptides were myristoylated at the N-terminal. All peptides were custom ordered from GenScript (Piscataway, NJ). 1mg of lyophilized peptides were initially dissolved in dimethyl sulfoxide (DMSO) to make a working stock solution. A final concentration of 200μM lipidated peptide was used when dissolved in PBS and the DMSO final concentration was less than 0.05%. Rats were first tested behaviorally for pain induction 4 days post-MIA prior to intraarticular peptide injection as previously described[19]. After confirmation of pain behavior, on day 5 rats were anesthetized with 2% isoflurane. The ipsilateral knee was sterilized, and an intraarticular injection of peptide was performed into the arthritic knee using a 28-guage X ½ inch needle with a polyethylene tubing allowing for a depth of 7 mm (to account for inflammation) was used for these injections and a Hamilton syringe. 50μL of lipidated peptide (200μM) was administered. The needle was kept in the knee joint for at least one minute to prevent solution leakage.

### 2.5 Dynamic Weight Bearing

Animals were given a 30-minute acclimation period in the testing room before initiating testing. Following habituation, animals were weighed and the weights were recorded. The animals were then placed within the dynamic weight bearing (DWB) chamber (BIOSEB, France). The chamber is connected to a weight sensor and a video camera allowing for automatic acquisition and scoring of weight behavior. Animals were given a 180 second latency period to explore the chamber, followed by a 300 second weight bearing acquisition period. Automatic acquisition data was validated manually for at least 90 seconds randomly using BIOSEB DWB-2 software to ensure accuracy of weight measurements.

### 2.6 von Frey testing

Animals were placed on top of an elevated wire-mesh platform (Ugo Basile, Italy) and allowed to acclimate to the testing chamber for 30 minutes. Touch Test Sensory Probes (Stoelting, Wood Dale, IL) were applied to the plantar surface of the ipsilateral and contralateral hindpaws. Filaments were applied in ascending or descending order using the Simplified-Up-Down method[6]. A middle filament of the series was presented to the animal, and a withdrawal response was recorded as either negative or positive. A negative response resulted in a larger filament size, while a positive response resulted in a filament of small size. Five filament presentations were given per animal with 5-minute intervals between filament presentations. The last filament response was used to calculate withdrawal threshold based on equations described previously[6].

### 2.7 Joint Histology

After completion of the MIA pain behavior assays on day 28, ipsilateral and contralateral knee joints were collected and fixed with 4% paraformaldehyde (PFA) for 48h at 4°C. Knee joints underwent decalcification by incubating them in 10% ethylenediaminetetraacetic acid for 14 days. Decalcified samples were washed in PBS and embedded in paraffin for sectioning. Sagittal sections (4μm) were placed on slides and stained with safranin-O. Sections were imaged using Aperio CS2 slide scanner (Leica Biosystems, Germany) at 20X magnification. Cartilage loss was scored according to the Osteoarthritis Research Society International (OARSI) scoring guidelines[3].

### 2.8 Micro-computed tomography

MicroCT imaging was performed at the Optical Imaging and Analysis Facility, School of Dental Medicine at the University at Buffalo as previously described[19]. After completion of the MIA pain behavior assay on day 28, ipsilateral and contralateral knee joints were collected and fixed with 4% (PFA) for 48h at 4°C and then washed with PBS. Ethanol (70%) was added to each sample, and each joint was placed in a ScanCo μCT100 scanner (Scanco Medical Brüttisellen, Switzerland) for imaging. The scanner parameters were set to 70 kVp for X-ray voltage and image resolution was kept at 10μm. Reconstructed DICOM files were imported to Image J software analysis (NIH). Tibial subchondral bone from the entire plateau was chosen as the region of interest for bone volume analysis. Bone volume was determined by assessing the bone volume fraction (BV/TV) using the Image J plugin BoneJ (NIH).

### 2.9 Statistics

All statistical analyses were performed using GraphPad Prism (GraphPad, La Jolla, CA). Comparisons between 2 groups were made using Student t test. Multiple comparisons were made using 2-way ANOVA followed by a Bonferroni post-hoc test. All data are presented as mean ± S.E.M. For area under the curve assessments, the curve was evaluated based on post-treatment trajectories and sustained improvement relative to the pre-treatment pain state, rather than absolute thresholds at a single pre-treatment time point.

## 3 Results

### 3.1 *In vivo* nociceptor AP2α2 knockdown attenuated OA pain-like behavior in mice

Male and female mice were given a single MIA injection unilaterally in the right knee to induce OA. To test the consequences of AP2α2 deficiency in nociceptors during MIA-induced OA, we ipsilaterally transfected the sciatic nerve with AP2α2-targeted shRNA after OA pain was established. We have previously shown that this *in vivo* shRNA transfection method results in a 60% reduction in AP2α2 protein after 7 days[15]. Mice received either AP2α2-targeted shRNA or control non-coding shRNA and tested behaviorally every 4 days for 28 days (Fig 1A). We observed an improvement in weight bearing 7 days after AP2α2 shRNA treatment, an improvement that lasted for about 19 days (Fig 1B). Similarly, we also observed a significant increase in paw withdrawal threshold after AP2α2 knockdown (Fig 1C). Data segregation based on sex can be found in Supplemental Figure 1. These data suggest that AP2α2 is necessary for pain processing in the MIA-OA pain model.

**Figure 1.**
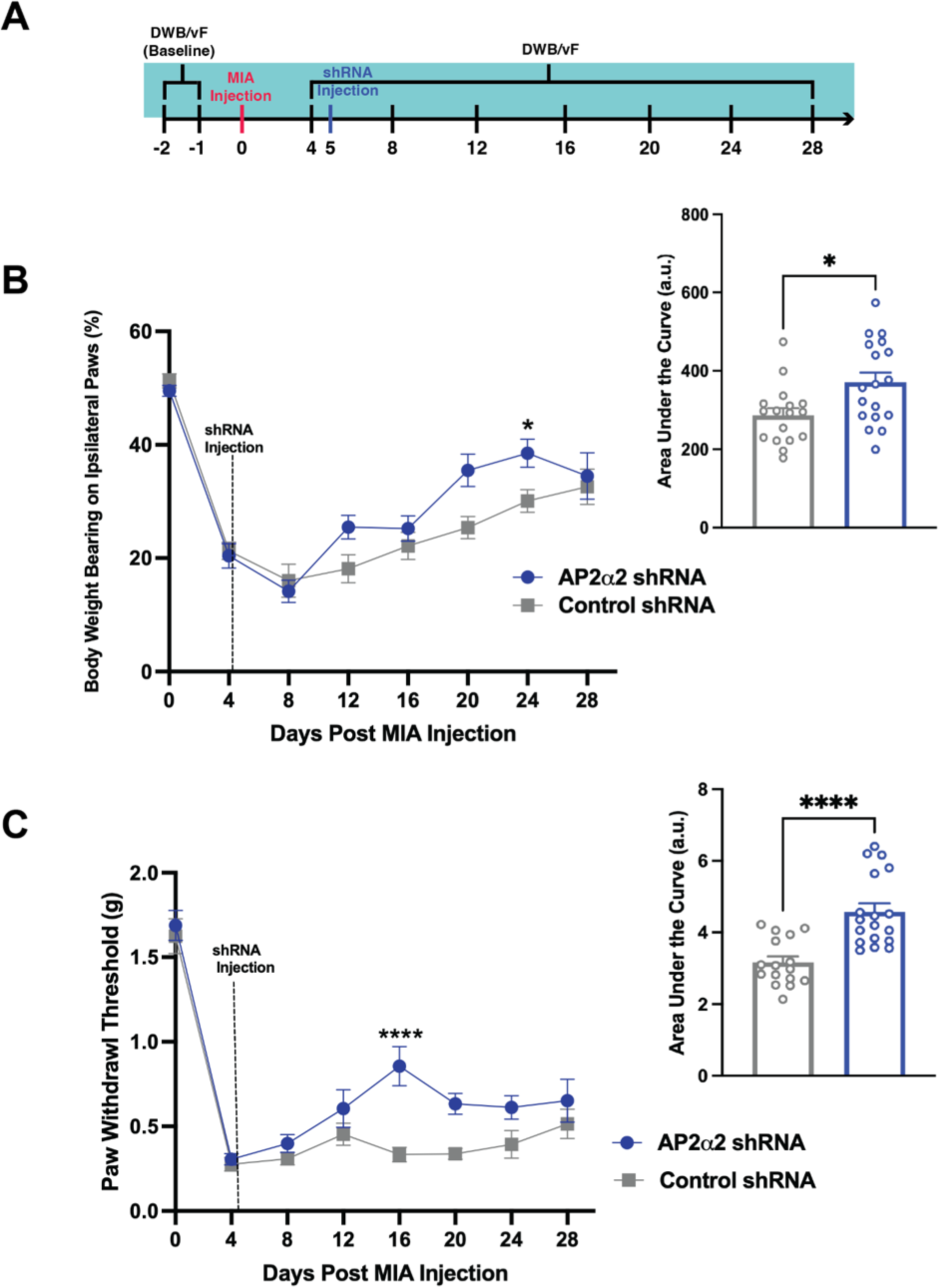
Genetic knockdown of AP2α2 attenuates pain-like behavior in mice. **A)** Experimental timeline for pain behavior assessment and injections of MIA and AP2α2 -targeted shRNA plasmid transfection. **B)** Percent of weight borne on the ipsilateral paw of mice post-MIA injection and AP2α2 or control shRNA plasmid. Pooled data for males and females is represented as cumulative mean ± S.E.M (Control n = 17, AP2α2 n=18). Significance determined by repeated measures 2-way ANOVA with Bonferroni correction; *p < 0.05. Total area under the curve of animals assessed for ipsilateral weight bearing. Significance determined by unpaired Student t-test *p < 0.05. **C)** von Frey withdrawal threshold (g) of ipsilateral paw of animals represented as cumulative mean ± S.E.M (Control n = 17, AP2α2 n=18). Significance determined by repeated measures 2-way ANOVA with Bonferroni correction; ****p < 0.0001. Total area under the curve for von Frey behavior. Significance determined by unpaired Student t-test; ****p < 0.0001.

### 3.2 Lipidated Ap2σ decoy peptide attenuated OA pain behavior in rats for multiple weeks after a single intraarticular injection

The AP2 adaptor complex uses two major endocytic motifs to bind to cargo proteins, one being an acidic di-leucine based motif which binds to the Ap2σ2 subunit[10]. We have previously demonstrated that a small, lipidated decoy peptide containing an acidic di-leucine motif, now referred to as A2σ inhibitor peptide, can prevent DRG neuronal hyperexcitability[4], and moderate acute inflammatory pain behaviors[15]. Here we tested whether a single intraarticular injection of the A2σ inhibitor peptide into an arthritic knee joint could similarly decrease OA pain behavior. The larger rat knee joint allowed for intraarticular injection[19]. Animals were given a single intraarticular injection of either Ap2σ inhibitor peptide or scrambled peptide (200μM, 50μL) 5 days after OA induction by MIA (Fig 2A). Pain behavior was assessed for 28 days after MIA. Ap2σ inhibitor peptide-treated animals showed significantly increased weight bearing on the arthritic ipsilateral side compared to scrambled peptide treated animals (Fig 2B).

**Figure 2.**
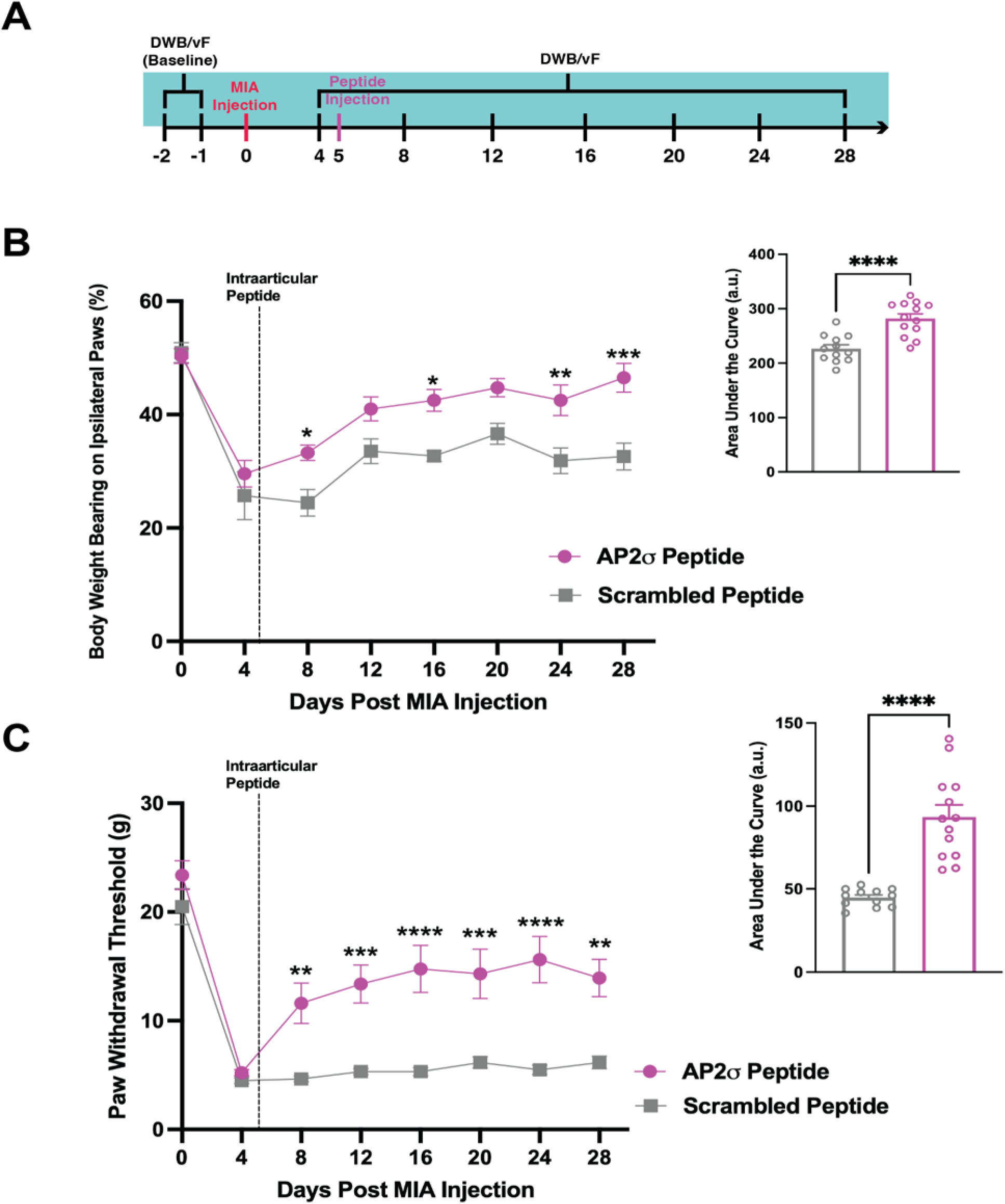
Single intra-articular injection of AP2σ inhibitor peptide attenuates OA pain-like behaviors in OA SD rats. **A)** Experimental timeline for pain behavior assessment and injections of MIA and AP2σ inhibitor or control peptide. **B)** Percent of weight borne on the ipsilateral paw post MIA injection and AP2σ inhibitor peptide or control peptide. Pooled data for males and females is represented as cumulative mean ± S.E.M (Control n = 12, AP2σ n=13). Significance determined by repeated measures 2-way ANOVA with Bonferroni correction; *p < 0.05; **p<0.001; ****p<0.0001 . Total area under the curve for animals assessed for ipsilateral weight bearing. Significance determined by unpaired Student t-test ****p<0.0001. **C)** von Frey withdrawal threshold (g) of ipsilateral paw of animals represented as cumulative mean ± S.E.M (Control n = 12, AP2α2 n=13). Significance determined by repeated measures 2-way ANOVA with Bonferroni correction; **p< 0.003; ***p<0.0005; ****p < 0.0001. Total area under the curve for von Frey behavior. Significance determined by unpaired Student t-test; ****p < 0.0001.

Mechanical sensitivity in the MIA assay is referred hyperalgesia where decreased thresholds reflect central sensitization[11]. Rats that received an intraarticular injection of the Ap2σ inhibitor peptide showed a significant increase in paw withdrawal threshold compared to scrambled peptide control (Fig 2C). In both weight bearing and mechanical sensitivity assessment, we observed significant reduction in pain behaviors that lasted throughout the duration of testing (Figs 2B&C). Data segregation based on sex is shown in Supplemental Figure 2. These data suggest that a single intraarticular injection of the peptide can reduce OA pain behavior for many weeks.

### 3.3 Histological confirmation of MIA-induced osteoarthritis in treated animals

To ensure that the MIA was producing cartilage loss, we collected knee joints from Ap2σ inhibitor peptide and scrambled peptide treated animals at the end of the 28-day behavioral assay. We determined cartilage integrity by staining with safranin-O and compared arthritic joints to the healthy non-MIA injected contralateral knee joints (Fig 3A). We observed significant cartilage loss in both Ap2σ inhibitor peptide and scrambled peptide treated animals (Fig 3B). Moreover, the cartilage loss was similar for Ap2σ inhibitor peptide and scrambled peptide demonstrating that Ap2σ inhibitor peptide-treated animals had arthritic knees.

**Figure 3.**
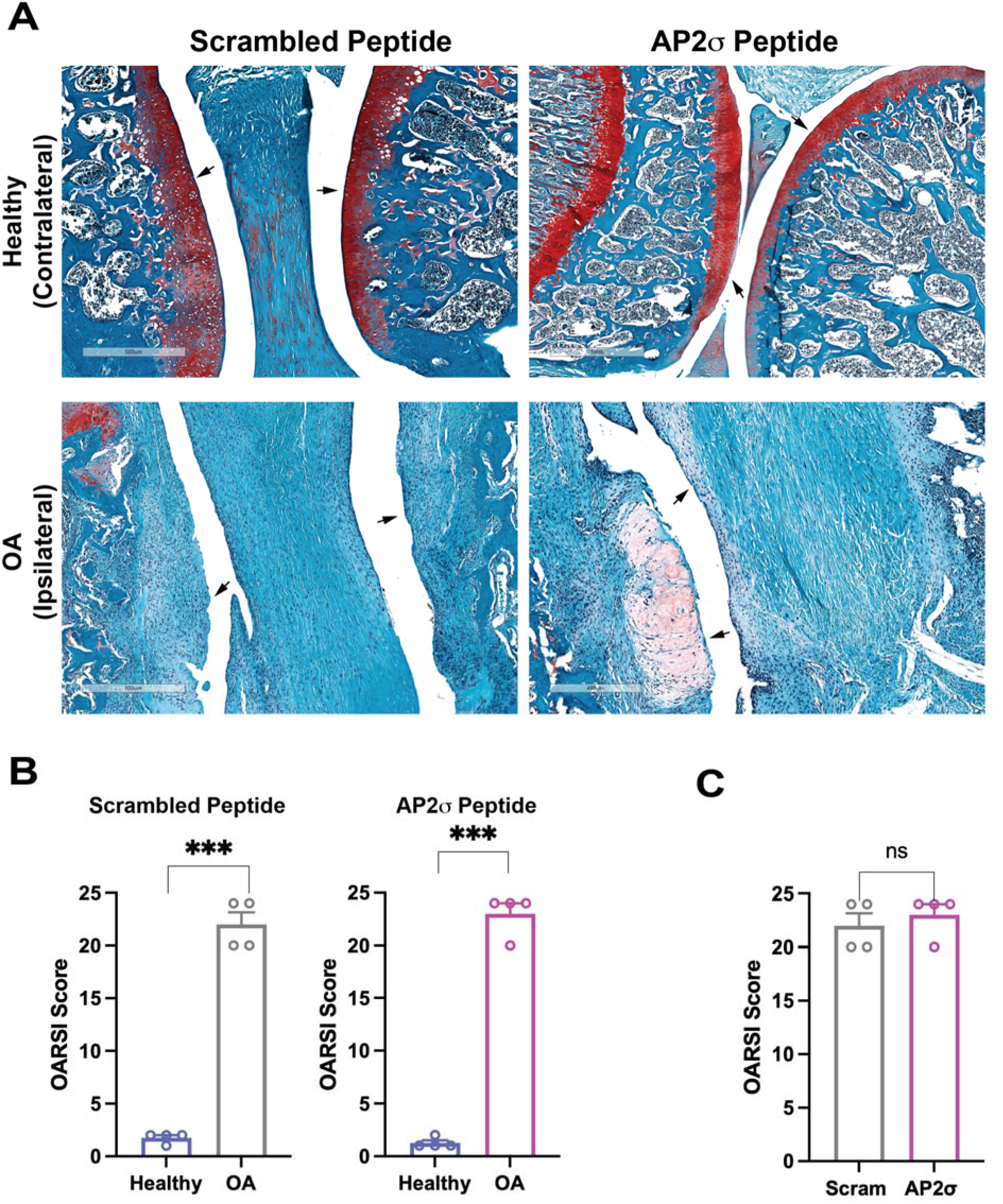
MIA-induced cartilage loss in treated groups. **A)** Representative images of tibial cartilage stained with Safranin-O 28 days post MIA injection. Samples received a single intra-articular injection of AP2σ inhibitor (n = 4) or scrambled peptide (n = 4) at day 4 post-MIA administration. Arrows indicate scored cartilage regions **B)** Cartilage integrity was determined via the OARSI scoring system and represented as cumulative means S.E.M. Ipsilateral AP2σ inhibitor- and scrambled peptide groups (OA) were compared to healthy contralateral groups. Significant differences were observed between scrambled and healthy contralateral knees. Significance was determined by paired Student t-test; ***p < 0.001. Significant differences were observed between AP2σ inhibitor and healthy contralateral knees. Significance was determined by paired Student t-test; ***p < 0.001. **C)** Comparison of AP2σ inhibitor vs. scrambled groups. Significance was evaluated using unpaired Student t-test. No significance was found.

### 3.4 Lipidated Ap2σ decoy peptide reduced OA-induced bone erosion

We also assessed joint integrity at the end of the MIA behavioral assay using micro-computed tomography (microCT). The MIA-OA model is characterized by erosion of subchondral bone[2; 24] with a significant reduction in bone volume fraction (bone volume/total volume: BV/TV%). A small lipidated peptide that induces the degradation of Na_V_1.8 channels [12; 17], when administered intraarticularly, effectively prevented bone loss in the MIA OA model as determined by microCT [19]. Here we similarly observed that while scrambled peptide-treated animals displayed significant bone loss 4 weeks after MIA (Figs 4A&B), Ap2σ inhibitor peptide-treated animals exhibited subchondral bone preservation (Fig 4A&C). Quantification of microCTdata revealed a statistically significant increase in BV/TV% in Ap2σ inhibitor peptide-treated animals compared to scrambled peptide-treated animals. These findings suggest that effective and sustained pain relief achieved through intraarticular Ap2σ inhibitor peptide administration is associated with a reduction in subchondral bone loss.

**Figure 4.**
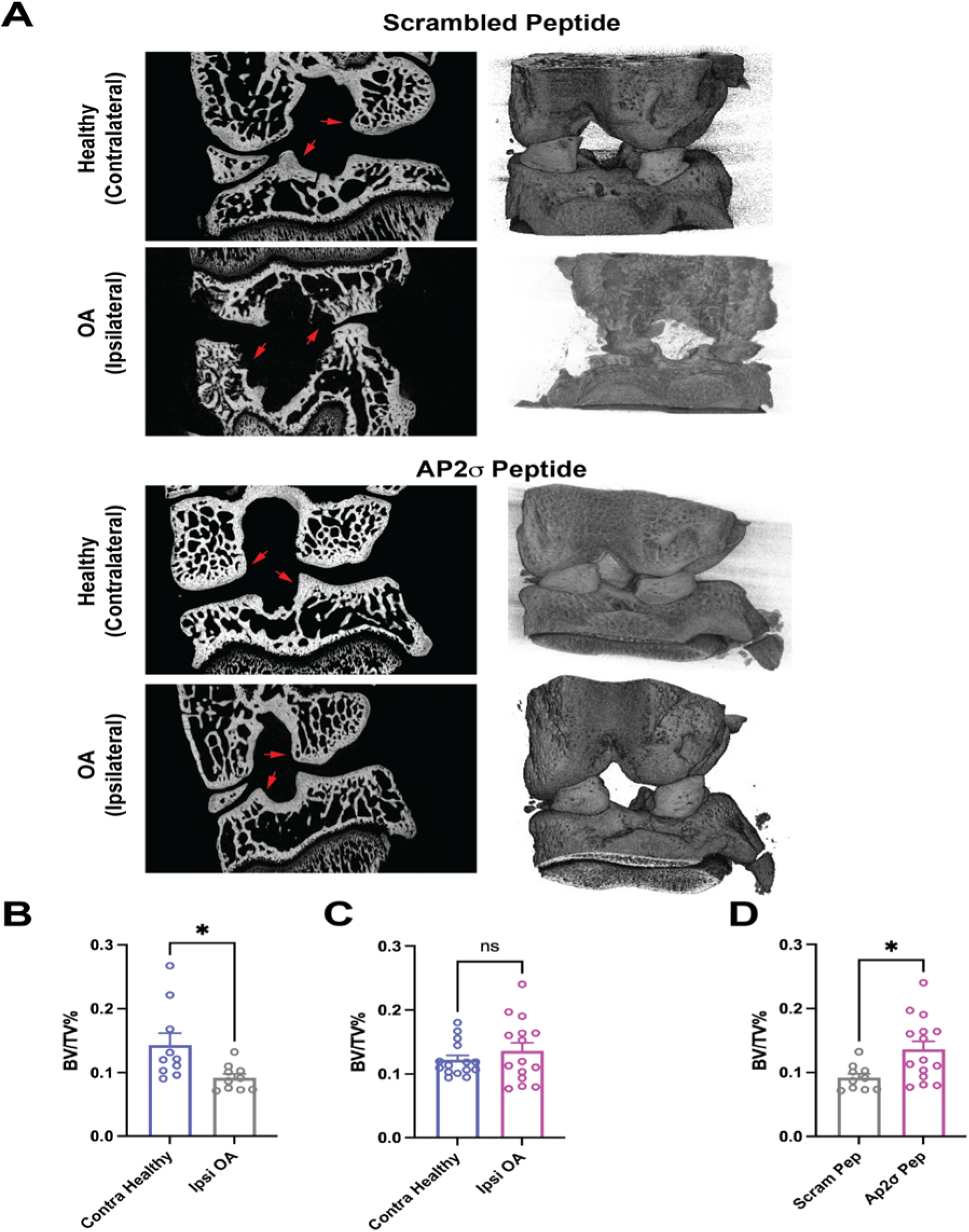
AP2σ inhibitor peptide treated animals exhibited reduced subchondral bone loss. **A)** Representative micro-computed tomography (microCT) images of rat knee subchondral bone 28 days after MIA-induced OA. Animals received a single intra-articular injection of scrambled peptide (n = 10) on day 4 post MIA administration. MicroCT images of knee subchondral bone of animals injected with AP2σ inhibitor peptide (n = 15) on day 4 post MIA administration. **B)** Quantification of bone volume fraction (BV/TV%) scrambled group (Healthy (Contralateral) vs OA (Ipsilateral)). **C)** Quantification of bone volume fraction (BV/TV%) AP2σ inhibitor peptide group (Healthy (Contralateral) vs OA (Ipsilateral)). Significance was determined by paired Student t-test; *p < 0.05. **D)** Comparison of AP2σ vs. scrambled groups. BV/TV analysis performed using ImageJ plugin BoneJ. Data was represented as cumulative means S.E. M. Significance was determined by unpaired Student t-test; *p < 0.05. BV/TV significant differences were observed in scrambled vs. healthy groups, and AP2σ vs. scrambled groups. No significant differences were detected in AP2σ vs. healthy groups.

## 4. Discussion

The AP-2 adaptor complex is multimeric protein consisting of 4 subunits: α, β2, μ2, and σ2. There are two genes that encode the α subunits: AP2α1 and AP2α2[18]. In neurons, AP2α1 concentrates at nerve terminals[1] and likely plays a prominent role in synaptic vesicle recycling while the AP2α2 is distributed more diffusely in cell bodies, axons and dendrites likely playing more of a role in recycling membrane components throughout the cell[1]. Within the dorsal root ganglia, AP2α2 is preferentially expressed in mouse and human CGRP^+^ nociceptors[15]. Importantly, the expression of AP2α2 was observed in peripherin-positive afferent terminals within human synovial tissue[9]. As we have demonstrated here, locally targeting the AP2 complex with a single intra-articular injection of the Ap2σ decoy peptide attenuated OA pain behavior for many weeks. Moreover, we demonstrated that this sustained pain reduction was accompanied by the preservation of subchondral bone, which is another indicator of the durable action of the Ap2σ inhibitor peptide.

In our previous studies, we employed the same genetic and pharmacological approach outlined herein to target Na_V_1.8 in the MIA-induced osteoarthritis (OA) model[19]. Within the knee joint, Na_V_1.8 expression is confined to nociceptors, whereas the AP2 complex is anticipated to exhibit a more extensive distribution, particularly in non-neuronal tissue. It was crucial for us to utilize nociceptor-specific genetic knockdown to reveal the contribution of nociceptor endocytosis to OA-induced pain. To achieve this we specifically targeted the AP2α2 gene using a previously validated mouse-specific AP2α2 shRNA[15] and an *in vivo* sciatic nerve knockdown strategy. Our findings revealed that AP2α2 deficiency in nociceptors resulted in a substantial increase in weight bearing and a reduction in mechanical sensitivity in mice (Fig 1). These data substantiated the notion that nociceptor endocytosis plays a significant role in pain signaling in OA, akin to the observations made in other models of inflammatory pain in both mice and rats[15].

We have previously demonstrated that the Ap2σ inhibitor peptide also reduced inflammatory pain behavior in both mice and rats[15]. To determine whether Ap2σ inhibitor peptide was efficacious in a non-recoverable model of inflammatory pain such as MIA-OA, it required the use of rats for their larger knees to conduct intraarticular injections[19]. We found a single intraarticular injection of the Ap2σ inhibitor peptide resulted in a significant increased weight bearing and decreased mechanical sensitivity for duration of the OA-pain behavior assay (Fig 2). This outcome contrasts with the findings for the PY(A) peptide, which targeted Na_V_1.8 channels, where the effect diminished after 19 days [19]. Several potential explanations exist for this observed disparity. Firstly, the Ap2σ inhibitor peptide specifically targets the acidic dileucine endocytotic motif, which has been demonstrated to inhibit the endocytosis of Slack K_Na_ channels[4]. This suggests that targeting Slack K_Na_ channels may be a more effective therapeutic approach than targeting Na_V_1.8 for the treatment of OA pain. Alternatively, the Ap2σ inhibitor peptide may exhibit greater potency than the PY(A) peptide. It is important to note that the concentrations employed in both studies (200μM at 50μL) are comparable. Nonetheless, we cannot negate the possibility that the Ap2σ inhibitor peptide is acting on both neuronal and non-neuronal tissue within the knee joint during OA. Importantly we have recently demonstrated that the Ap2σ inhibitor peptide did not block all endocytosis (Nicosia and Bhattacharjee 2026 in press) as it failed to block transferrin uptake in cells which is controlled by the YxxΦ endocytotic motif and the AP2μ subunit. Still, it is possible that Ap2σ inhibitor peptide is affecting resident inflammatory cells within the arthritic joint. Although we did not observe changes in cartilage destruction in Ap2σ inhibitor peptide-treated animals (Fig 3), it is possible that pain signaling might have been affected by the effect of the Ap2σ inhibitor peptide on non-neuronal cells. These possibilities will require further exploration.

We observed that the Ap2σ inhibitor peptide mitigated MIA-induced subchondral bone erosion. In this case, the Ap2σ inhibitor peptide acted similarly to the PY(A) peptide targeting Na_V_1.8 channels [19]. A single intraarticular injection of Ap2σ inhibitor peptide resulted in the preservation of subchondral bone 24 days after MIA injection (Fig 4). Although bone remodeling in osteoarthritis (OA) is complex, early OA is characterized by bone loss[25]. While the MIA model cannot capture all aspects of the arthritic disease, subchondral bone loss does seem to be a feature of this model[2; 24] As we described before, mechanical forces are crucial for bone homeostasis[24]. The effective and sustained reduction in pain produced by the Ap2σ inhibitor peptide and the concomitant increased weight bearing (Fig 2) likely contributed to the reduction in MIA-induced bone loss. As we have noted previously, for the MIA-OA pain behavior assay specifically, microCT analysis of subchondral bone may serve as an additional measure of effective and prolonged analgesia.

Intraarticular interventions are gradually assuming a more significant role in the treatment of osteoarthritis, although their long-term structural effects remain uncertain [20]. A meta-analysis of various orthobiologics, including platelet-rich plasma, stromal vascular fraction, and bone marrow aspirate concentrate, failed to demonstrate clinically meaningful differences in pain scores compared to viscosupplementation[5]. Most intraarticular injectables, such as corticosteroids, primarily target inflammation rather than nociceptors directly. The results presented herein with the Ap2σ inhibitor peptide, as well as those previously obtained with the PY(A) peptide[19] suggest that small lipidated peptides targeting nociceptors may represent a novel approach for the treatment of osteoarthritis pain.

## Acknowledgments

The authors would like to thank Dr. Imtiaz Mohammed of the University at Buffalo Histology Core facility for his assistance with knee joint safranin-O staining. We would also like to thank Dr. Andrew McCall for his assistance with MicroCT scanning at the Optical Imaging and Analysis facility.

## Author Contributions

AB and AJC conceived the idea for the project and co-wrote the manuscript. AJC performed all the behavioral, histology, knee imaging and immunohistochemistry experiments, analyzed data, and generated the manuscript. JST piloted initial MIA studies in rats. RR performed initial analysis on microCT and bone volume.

