## Supplemental Figure for "Inhibiting nociceptor endocytosis reduces MIA-induced osteoarthritic pain behavior"

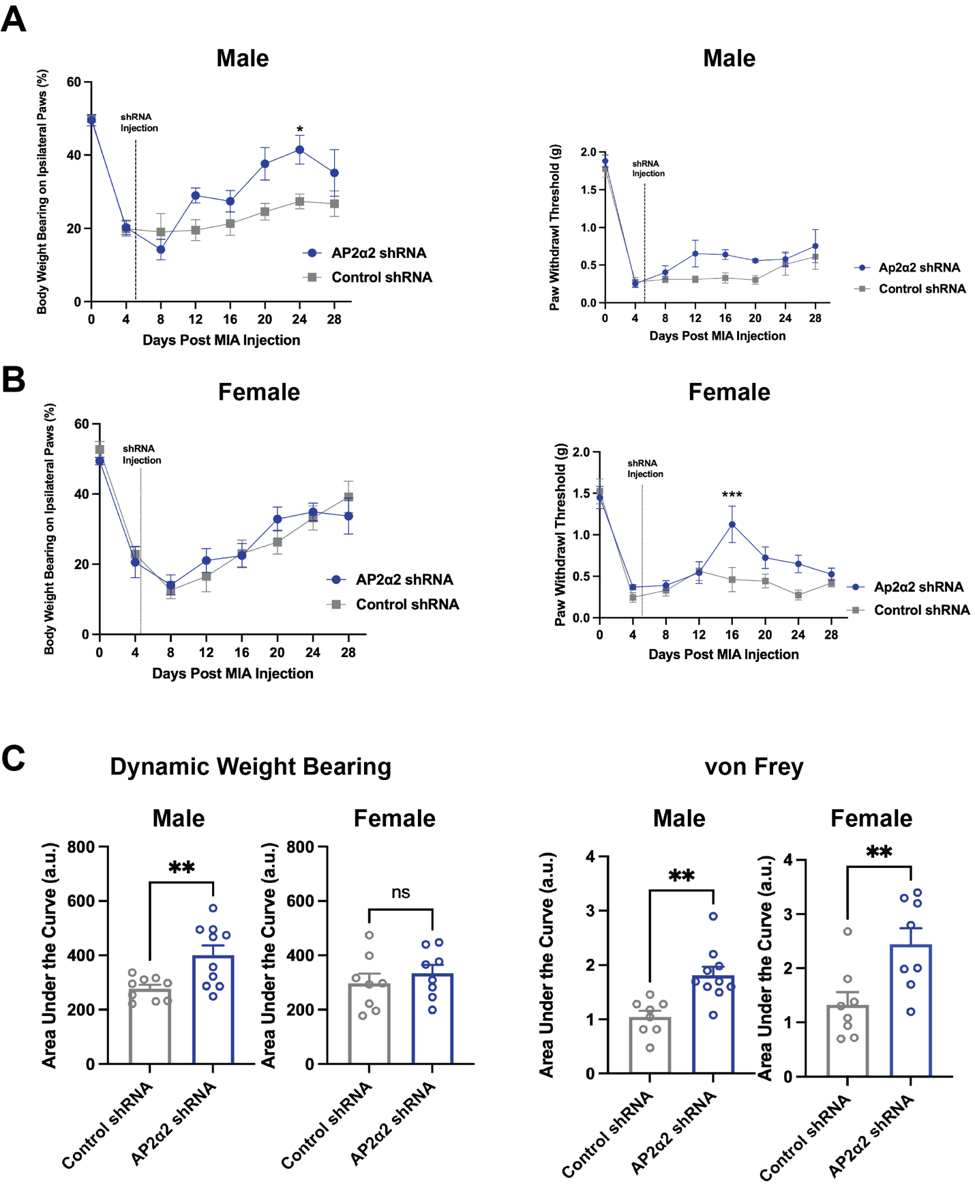

**Supplemental Figure 1. AP2α2 knockdown attenuates pain-like behavior in OA mice segregated by sex.**

C57BL/6 mice were used to test the effects of a sciatic nerve transfection of AP2α2 targeted shRNA plasmid upon OA pain. Transfection occurred 4 days post MIA injection within the knee joint and pain behavior was recorded at baseline, 4 days post MIA, and every four days thereafter until completion of the experiments on day 28. **A)** Male mice were assigned to either the AP2α2 shRNA plasmid group (n=8) or the control shRNA group (n=8) Percent of weight borne on the ipsilateral paw of animals after MIA injection and AP2α2 or control shRNA plasmid. Data for males is represented as cumulative mean ± S.E.M (n = 7 per group). Significance determined by repeated measures 2-way ANOVA with Bonferroni correction *p < 0.05. von Frey withdrawal threshold (g) of ipsilateral paw of animals represented as cumulative mean ± S.E.M. Significance determined by repeated measures 2-way ANOVA with Bonferroni correction. **B)** Female mice were assigned to either the AP2α2 shRNA plasmid group (n=8) or the control shRNA group (n=8). Percent of weight borne on the ipsilateral paw of animals after MIA injection and AP2α2 or control shRNA plasmid. Data for females is represented as cumulative mean ± S.E.M (n = 7 per group). Significance determined by repeated measures 2-way ANOVA with Bonferroni correction. von Frey withdrawal threshold (g) of ipsilateral paw of animals represented as cumulative mean ± S.E.M. Significance determined by repeated measures 2-way ANOVA with Bonferroni correction ***p < 0.001. **C)** Total area under the curve for animals assessed for ipsilateral weight bearing behavior. Significance determined by unpaired Student t test. For males **p<0.001. For females significance was not found. Total area under the curve for von Frey behavior. Significance determined by unpaired Student t test. For males **p<0.001. For females **p<0.001.

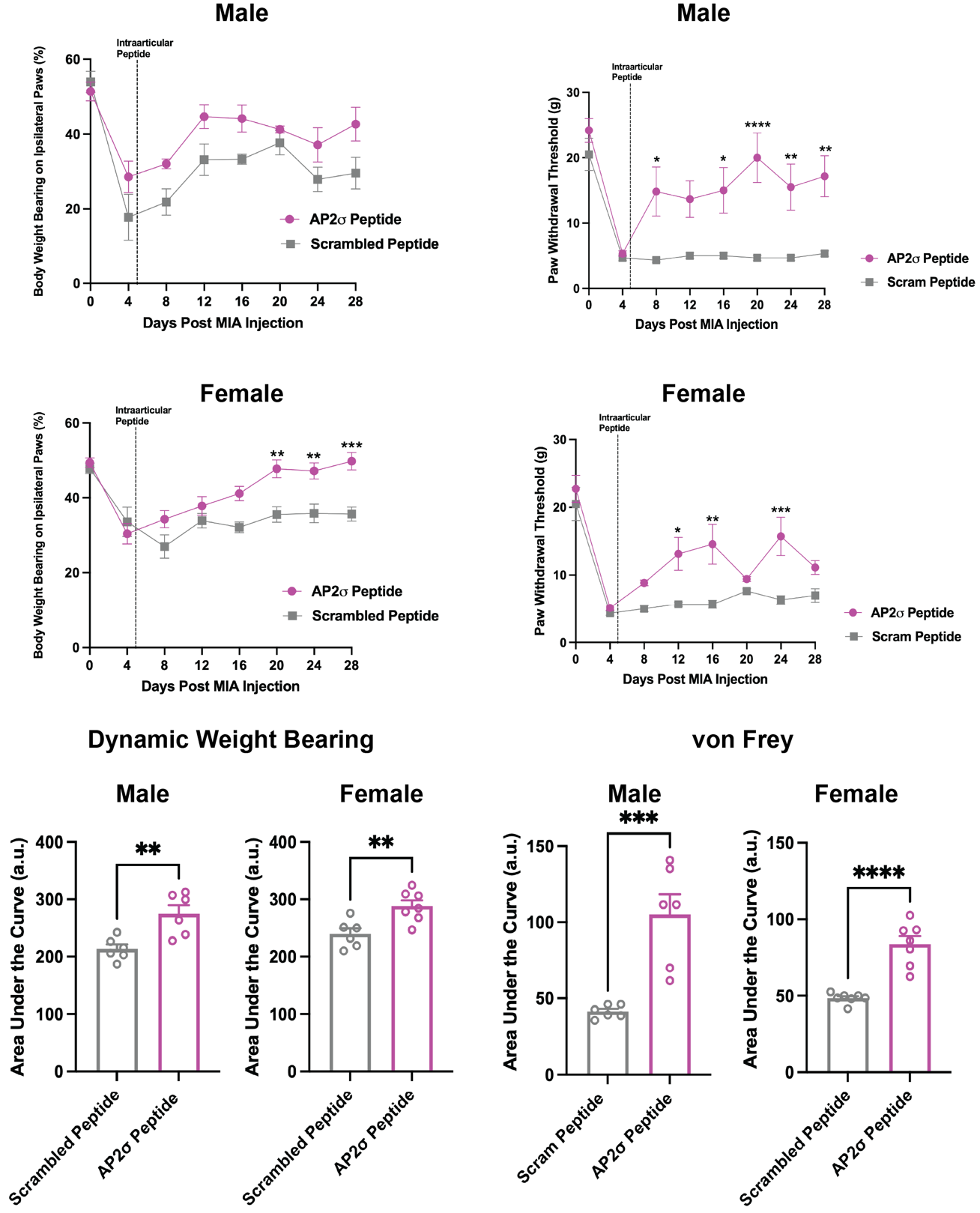

**A)** Males: MIA was injected intraarticularly (IA) 4 days before AP2σ inhibitor peptide (n=6) or scrambled peptide (n=6) was injected IA. Weight bearing was measured as previously stated. Data for males are represented as cumulative mean ± S.E.M. Significance determined by repeated measures 2-way ANOVA with Bonferroni correction p < 0.05; *p < 0.01; **p < 0.001; *** (AP2σ vs. Scrambled). von Frey withdrawal threshold (g) of ipsilateral paw of animals represented as cumulative mean ± S.E.M. Significance determined by repeated measures 2-way ANOVA with Bonferroni correction *p < 0.01; **p < 0.001; ***p<0.0001; ****p<0.00001. **B)** Females: MIA was injected intraarticularly (IA) 4 days before AP2σ inhibitor peptide (n=7) or scrambled peptide (n=6) was injected IA. Dynamic weight bearing data for females are represented as cumulative mean ± S.E.M. Significance determined by repeated measures 2-way ANOVA with Bonferroni correction **p < 0.01; ***p < 0.001 (AP2σ vs. scrambled peptide). von Frey withdrawal threshold data represented as cumulative mean ± S.E.M. Significance determined by repeated measures 2-way ANOVA with Bonferroni correction *p<0.01; **p<0.001; ***p < 0.001. **C)** Total area under the curve of animals assessed for ipsilateral weight bearing behavior. Significance determined by unpaired Student t test. For males **p<0.001. For females **p<0.001. Total area under the curve for von Frey behavior. Significance determined by unpaired Student t test. For males ***p<0.0001. For females ****p<0.00001.

**Supplemental Figure 2. Injection of a lipidated AP2σ inhibitor peptide attenuates pain-like behavior in OA rats segregated by sex**
